# Sex Differences in Solute and Water Handling in the Human Kidney: Modeling and Functional Implications

**DOI:** 10.1101/2021.02.03.429526

**Authors:** Rui Hu, Alicia A. McDonough, Anita T. Layton

## Abstract

Besides the excretion of metabolic wastes, the kidneys regulate homeostasis of electrolytes, pH, metabolites, volume and blood pressure. Sex differences in kidney function and blood pressure have been widely described across many species. Immunoblot analysis has revealed that the kidney of a female rat is not simply a smaller version of a male kidney. Rather, male and female rat kidneys exhibit dimorphic patterns of transporter expression and salt handling, the functional implications of which have been analyzed in a series of previously published modeling studies of rat kidney function. In the present study, we extend the analysis to the human kidney: we developed sex-specific models of solute and water transport in the human kidney, and identified epithelial transport parameters, consistent with patterns found in male and female rats, that yield urine output and excretion rates consistent with known human values. The model predicts that the lower sodium hydrogen exchanger 3 (NHE3) activity in women reduces the fractional reabsorption of Na^+^, K^+^, Cl^-^, and water along the proximal tubule, compared to men, and that the larger load on the distal nephron can be handled by enhanced activities in key Na^+^ transporter such as epithelial sodium channel (ENaC) and sodium chloride cotransporter (NCC) in women. Model simulations further indicate that the larger distal transport capacity and proximal transport reserve may better prepare women for elevated demands of pregnancy and lactation. The larger distal transport capacity may also contribute to reduced efficacy of angiotensin converting enzyme inhibitors to lower blood pressure in women.

**Author summary:** The kidneys maintain homeostasis by controlling the amount of water, ions, and other substances in the blood. That function is accomplished by the nephrons, which transform glomerular filtrate into urine by an exquisite transport process mediated by a number of membrane transporters. Recently, the distribution of renal transporters along the nephron has been shown to be markedly different between male and female rodents. We postulate that similar sexual dimorphism exists between men and women, and we seek to reveal its physiological implications. We hypothesize that the larger abundance of a renal Na^+^ transport in the proximal tubules in females may also better prepare them for the fluid retention adaptations required during pregnancy and lactation, durint which renal and systemic hemodynamics are both drastically altered by the marked volume expansion and vasodilation. Also, kidneys play a key role in blood pressure regulation, and a popular class of anti-hypertensive medications, angiotensin converting enzymes (ACE) inhibitors, have been reported to be less effective in women. Model simulations suggest that the blunted natriuretic and diuretic effects of ACE inhibition in women can be attributed, in part, to their higher distal baseline transport capacity.

## Introduction

Throughout the animal kingdom kidneys are known primarily for their function as filters, removing metabolic wastes and toxins from the blood for excretion in the urine. But in mammals, kidneys specialized to serve various other essential regulatory functions, including water, electrolyte and acid-base balance [1]. In humans, the pair of kidneys are located in the abdominal cavity, with one on each side of the spine. A mammalian kidney can be divided into an outer region (cortex) and an inner region (medulla). Each of the human kidneys contains about a million glomeruli, which are clusters of capillaries that each receive blood from individual afferent arterioles branching off intra-renal arteries. Driven by vascular hydrostatic pressure, a fraction of the water and solutes in that blood is filtered through the glomerulus and becomes the tubular fluid of the nephron. The nephrons adjust the glomerular filtrate, via absorptive and secretive processes, mediated by membrane transporters and channels on the renal tubular epithelial cells. Thus, tubular fluid begins at the glomerulus as an ultrafiltrate of plasma and is transformed into urine at the end of the nephrons [1]. Regulation of the epithelial transport processes that match urine output to both intake of fluids and solutes as well as to waste product production is the subject of a large body of experimental and theoretical effort [2–4]. Computational models have been developed to unravel the renal solute and water transport processes in humans [5] and rats, under dietary [6, 7] and therapeutic [8–10] manipulations, and under pathophysiological conditions [11, 12].

In recent years, a new dimension for investigation has emerged: sex. Like nearly every tissue and organ in the mammalian body, the structure and function of the kidney is regulated by sex hormones [13–17]. Veiras et al. [18] reported sexually dimorphic patterns in the abundance of electrolyte transporters, channels, and claudins (collectively referred to as transporters) in male and female rodents. Their findings demonstrated that, compared with male rat nephrons, female rat nephrons exhibit lower activities of major Na^+^ and water transporters along the proximal portion of the renal tubule (proximal tubule), resulting in significantly larger fractional delivery of Na^+^ and water to the downstream nephron segments in female kidneys. Along the distal nephron segments, the female kidney exhibits a higher abundance of key Na^+^ transporters, relative to male, resulting in similar urine excretion between the sexes. To assess the functional implications of these findings, our group developed sex-specific rat nephron transport models [19–21]. Simulation results suggest an explanation for the more rapid excretion of a saline load in female rats observed in Ref. [18]: In response to a saline load, the Na^+^ load delivered distally is substantially greater in female rats than males, overwhelming transport capacity, cumulating in more rapid compensatory natriuresis in female rats.

Some of these sex differences in rodent kidney function may translate to humans. Across race, men have higher blood pressure than women through much of lifespan [22, 23]. Given the essential role of the kidney in blood pressure control, sex differences in hypertension may be attributable, in part, to differences in kidney structure and function. That said, it is noteworthy that some significant differences in renal transporter patterns were reported between rats and mice [18], and the differences between rat and human kidneys may be greater. For example, in rats, male kidneys are bigger than female kidneys, whereas men and women have kidneys of similar size [24]. Nonetheless, like female rats, women also face the challenge of circulating volume adaptation during pregnancy and lactation, and the potential for increasing reabsorption along the proximal nephron, coupled to the more abundant transporters in the distal nephron, that may facilitate these adaptations. Given these observations, the objective of this study is to answer the questions: 1) To what extent can observed differences in blood pressure regulation between men and women be accounted for by sexual dimorphism in renal transporter patterns? 2) How do these differences in renal transporter pattern permit the reserve renal capacity that allows women to better handle the glomerular hyperfiltration that they face in pregnancy?

## Results

### Sex differences in segmental transport under baseline conditions

The amounts of solutes and water reabsorbed or secreted along a given nephron segment depends, in part, on the activity of membrane transporters. We assume that human nephrons exhibit sexual dimorphism in membrane transporter patterns (density) similar to that observed in rats [18]. Unlike rats and mice, kidneys in men and women have similar size [24], thus, similar effective tubular transport areas, as well as similar glomerular filtration rate (GFR) and urine output. Given the similarities in size and filtration between the human sexes, to what extent might sex differences in renal transporter expressions alter segmental solute and transport patterns? To answer that question, we conduct model simulations under control conditions and compare solute and water transport along individual segments in men and women.

Figure 2 shows the predicted delivery of key solutes [Na^+^, K^+^, Cl^-^, HCO_3_^-^, NH_4_^+^, urea, and titratable acid (TA)] and fluid to the inlets of individual superficial and juxtamedullary nephron segments, expressed per kidney, in the male and female models. [TA] is computed from [H_2_PO_4_^+^] and [HPO_4_^2-^]. Single-nephron GFR (SNGFR) is assumed to be 100 and 133.3 nl/min in the superficial and juxtamedullary nephrons, respectively, in both sexes. Recall also that there are almost 6 times as many superficial than juxtamedullary nephrons, thus the smaller juxtamedullary contribution per kidney despite the higher SNGFR in the juxtamedullary nephrons (see Fig. 2). Solute and fluid transport rates along individual superficial and juxtamedullary nephron segments are shown in Fig. 3. Tubular fluid solute concentrations and luminal fluid flow are shown in Figs. S1 and S2 in supporting information.

**Figure 1.**
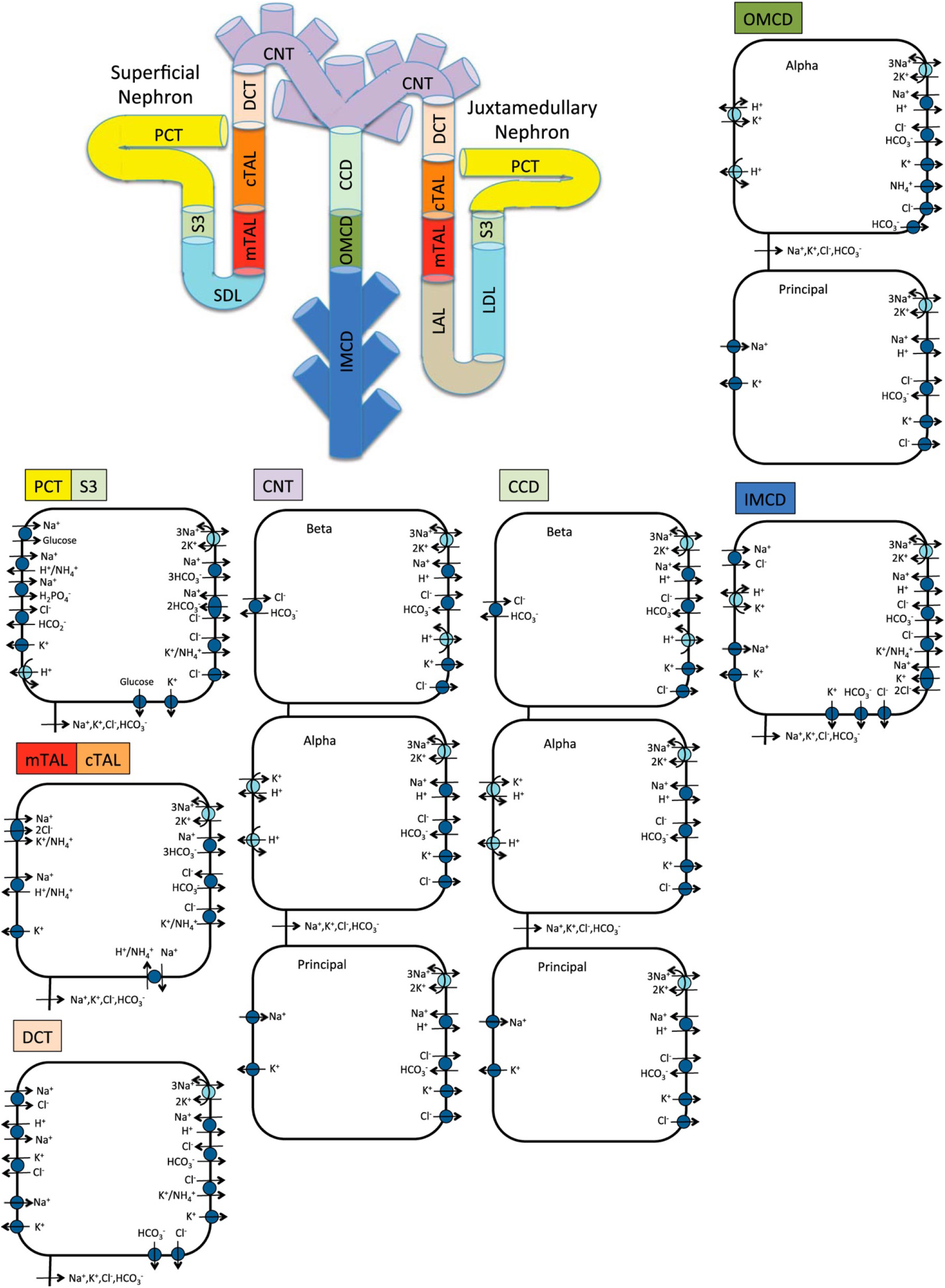
Schematic diagram of the nephron system (not to scale). The model includes 1 representative superficial nephron and 5 representative juxtamedullary nephrons, each scaled by the appropriate population ratio. Only the superficial nephron and one juxtamedullary nephron are shown. Along each nephron, the model accounts for the transport of water and 15 solutes (see text). The diagram displays only the main Na^+^, K^+^, and Cl^-^ transporters. Red boxes highlight model membrane transporters whose expressions differ between males and females. mTAL, medullary thick ascending limb; cTAL, cortical thick ascending limb; DCT, distal convoluted tubule; PCT, proximal convoluted tubule; CNT, connecting duct; CCD, cortical collecting duct; SDL, short or outer-medullary descending limb; LDL/LAL, thin descending/ascending limb; OMCD, outer-medullary collecting duct; IMCD, inner-medullary collecting duct.

**Figure 2.**
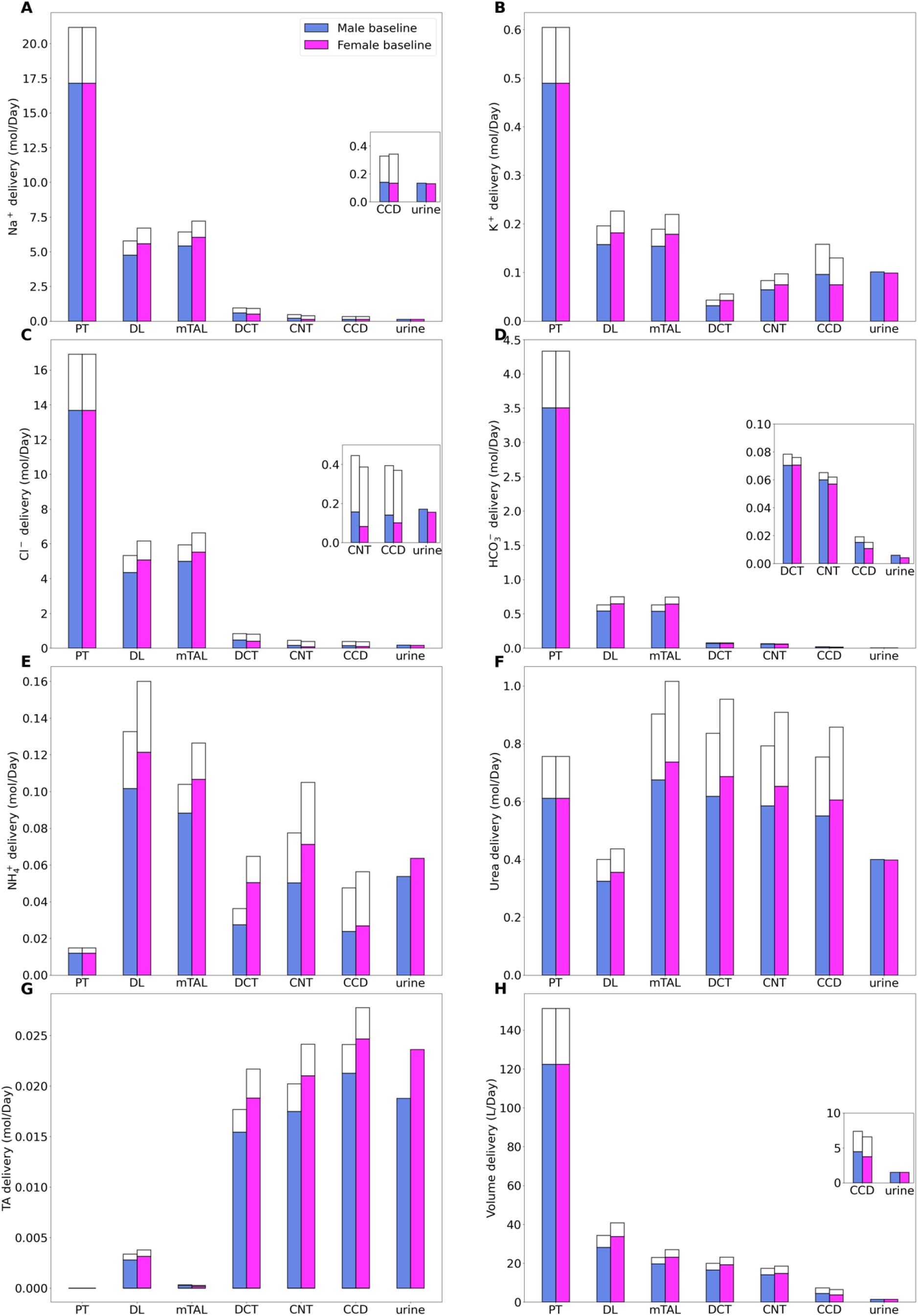
Delivery of key solutes (*A*–*G*) and fluid *(H)* to the beginning of individual nephron segments in men and women. Color bars, superficial nephron values; white bars, juxtamedullary values, computed as weighed totals of the five representative model juxtamedullary nephrons. The model assumes a superficial-to-juxtamedullary nephron ratio of 85:15, thus, the superficial delivery values are generally higher. In each panel, the two bars for “urine” are identical since the superficial and juxtamedullary nephrons have merged at the cortical collecting duct entrance. PT, proximal tubule; DL, descending limb; mTAL, medullary thick ascending limb; DCT, distal convoluted tubule; CNT, connecting tubule; CCD, cortical collecting duct; TA, titratable acid. *Insets:* reproductions of distal segment values.

**Figure 3.**
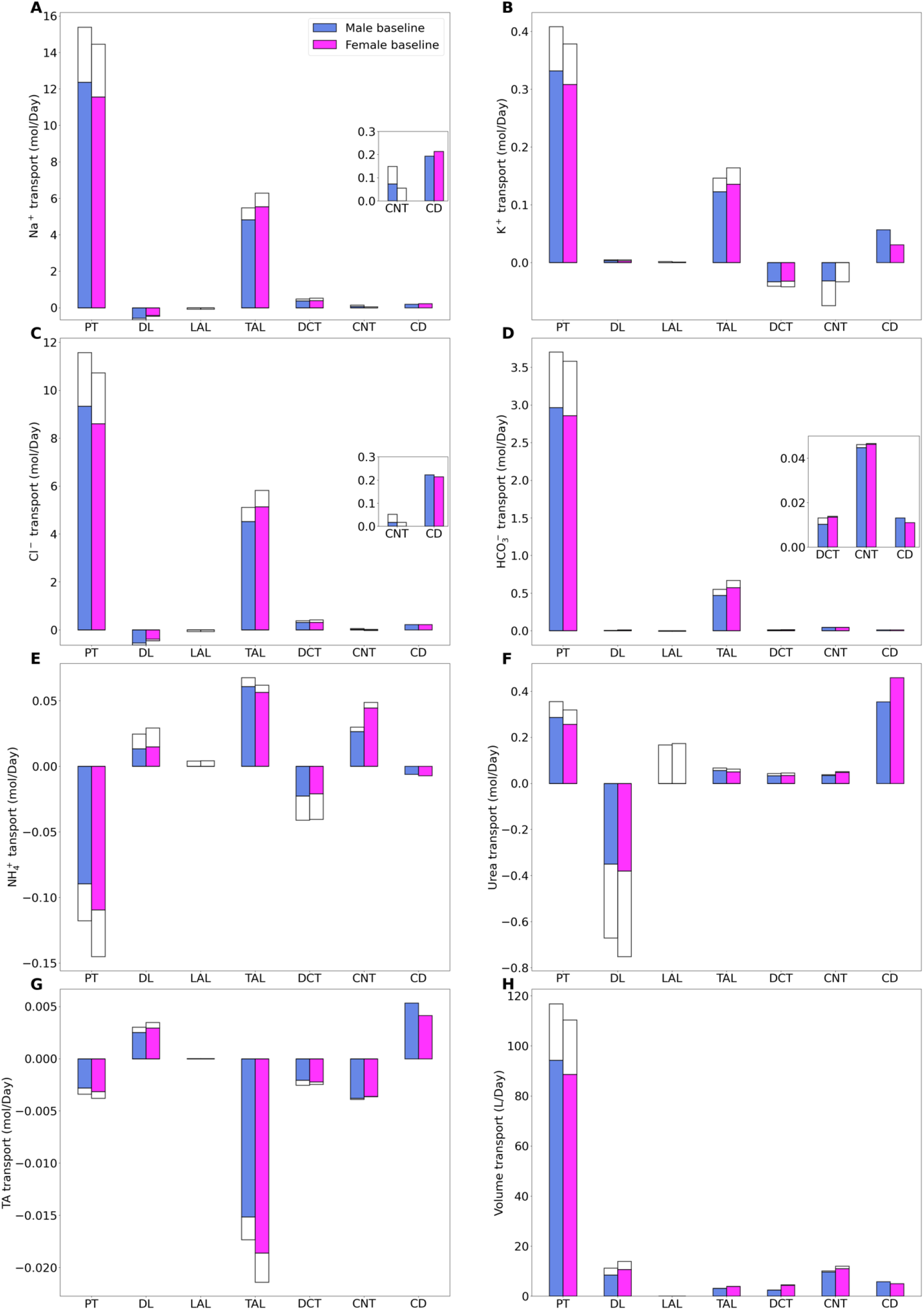
Net transport of key solutes (A–G) and fluid (H) along individual nephron segments, in men and women. Transport is taken positive out of a nephron segment. PT, proximal tubule; DL, descending limb; TAL, thick ascending limb; DCT, distal convoluted tubule; CNT, connecting tubule; CD, collecting duct; TA, titratable acid. Insets: reproductions of distal segment values. Notations are analogous to Fig. 2.

In both men and women, the majority of the filtered Na^+^ is reabsorbed by the proximal tubules. In men, fractional Na^+^ reabsorption is predicted to be 73%; in women, where the activity of the principal Na^+^ transporter sodium hydrogen exchanger 3 (NHE3) is assumed to be 83% of men, 68%. Most of the remaining Na^+^ is reabsorbed by the thick ascending limbs. With Na^+^ delivery to the thick ascending limbs higher in women, we hypothesize that Na-K-Cl cotransporter (NKCC) activity is substantially higher in women (Table 1). Na-Cl cotransport (NCC) has been shown to be higher in women [25]. With this configuration, Na^+^ flows along the distal tubular segments are similar between men and women, as is urine Na^+^ excretion, at ~0.6% of filtered Na^+^ (Figs. 2A and 3A). The transport of Cl^-^, the accompanying anion, follows trends similar to Na^+^ transport (Figs. 2C and 3C).

**Table 1.**
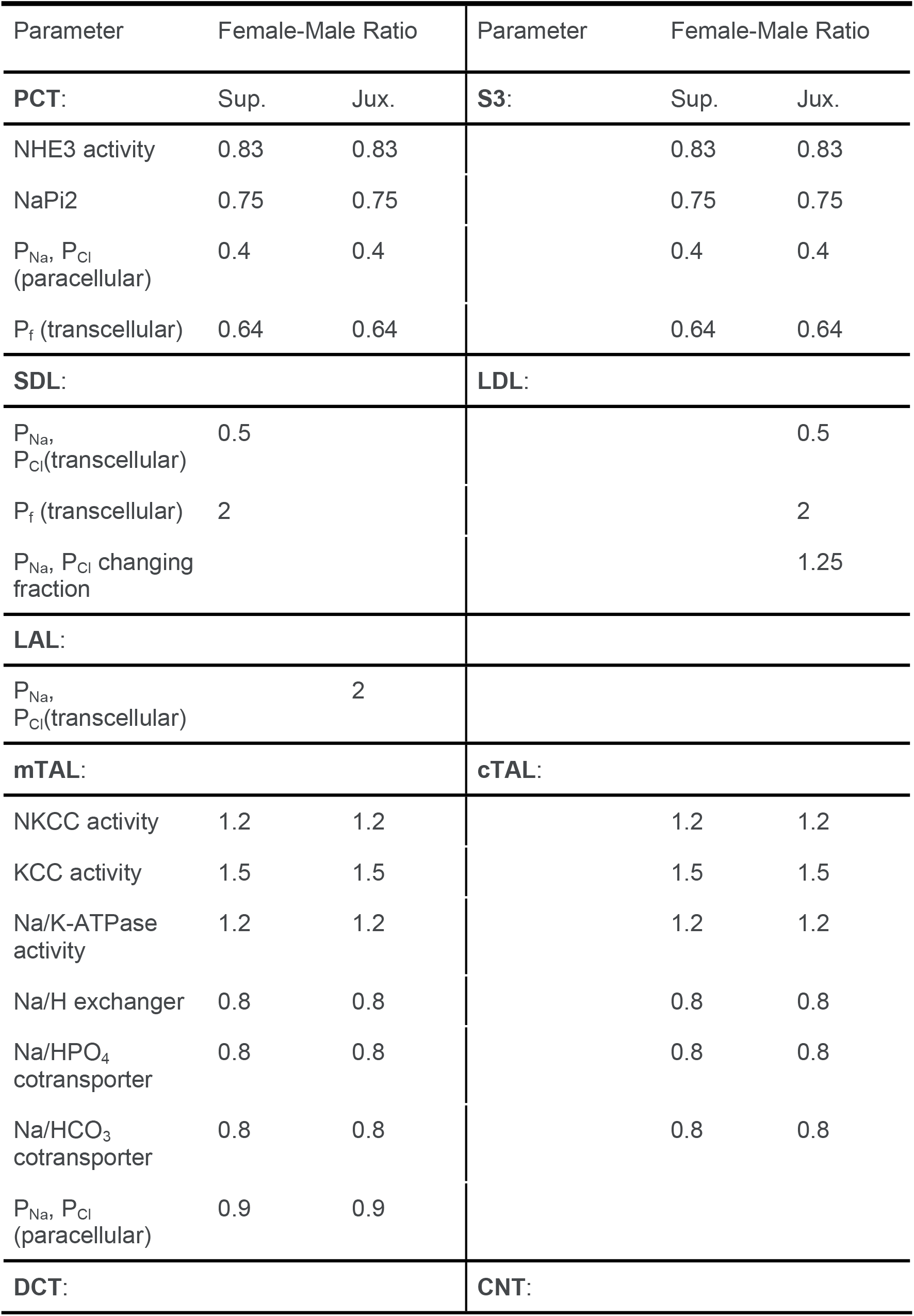
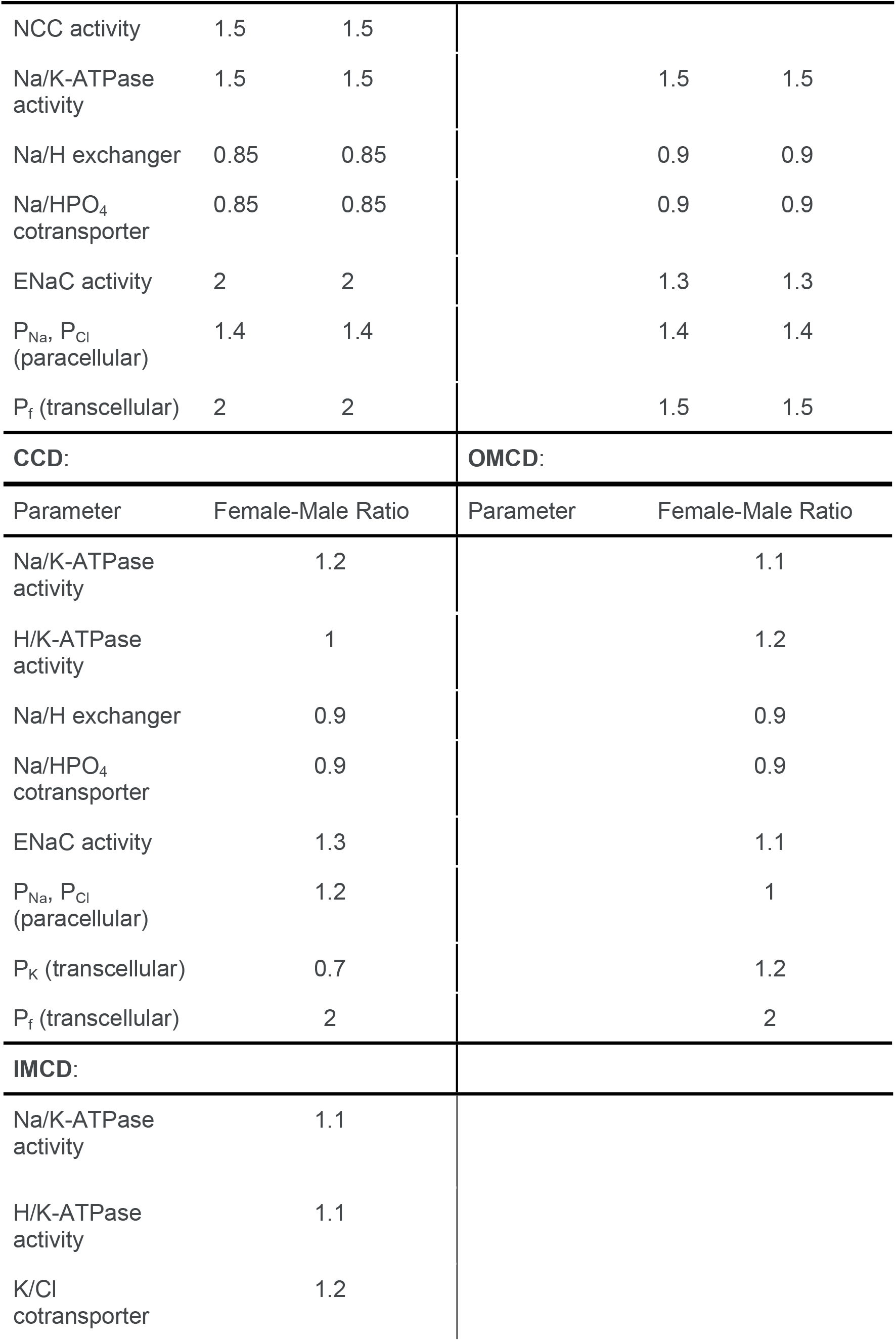
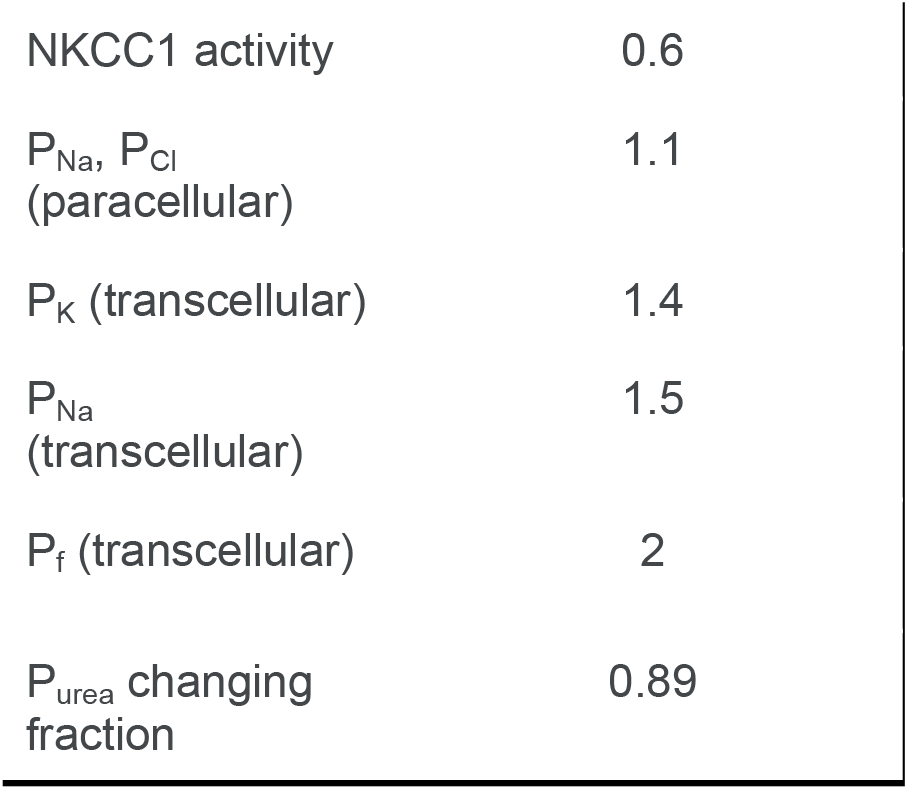
Key differences in nephron transport parameters between men and women.

Water transport is driven principally by the osmotic gradient; as such, along nephron segments that have sufficiently high water permeabilities, Na^+^ reabsorption is accompanied by water reabsorption. The model predicts that 77% of the filtered volume is reabsorbed along the proximal tubules in men and 73% in women (Figs. 2H and 3H). Because the thick ascending limbs have low water permeability, much of the Na^+^ reabsorption along that segment is unaccompanied by water, resulting in a precipitous drop in tubular fluid osmolality from 700 at the inlet to 300 mosm/(kg H_2_O) at the outflow of the thick ascending limbs. That substantial transcellular osmolality gradient, together with additional NaCl transport, drives water reabsorption along the water-permeable segments of the distal nephron. The model predicts that ~1% of the filtered volume is excreted.

In terms of filtered K^+^ loads, the model predicts that in men 68% of the filtered K^+^ is reabsorbed along the proximal tubules, slightly higher than the 63% in women. The majority of the remaining K^+^ is reabsorbed along the thick ascending limbs. Downstream of the Henle’s loops, the model connecting tubules are assumed to vigorously secrete K^+^. Due to their longer length, juxtamedullary connecting tubules secrete substantially more K^+^ than the superficial segments. Model parameters were chosen so that a similar fraction of the filtered K^+^ load (16-17%) is excreted in men and women. See Figs. 2B and 3B.

In both sexes, substantial NH_4_^+^ secretion occurs along the proximal tubules, via substitution of H^+^ in the NHE3 transporter and secretion of NH_3_. Downstream along the thick ascending limbs, a substantial fraction of NH_4_^+^ is reabsorbed substituting for K^+^ in NKCC2. NH_4_^+^ excretion is predicted to be 19% higher in women (Fig. 2E), consistent with the higher ammonia excretion reported in female mice, despite similar protein intake as males [26]. Urea is reabsorbed along most nephron segments, except the descending limbs of the loops of Henle, where urea enters due to the presumed higher interstitial urea concentration (Fig. 2F).

### Female pattern provides renal transport reserve capacity necessary in to preserve water and salt in challenges of reproduction and lactation

The baseline simulations indicate that, given the same renal hemodynamics (input), both the male and female renal transporter patterns can yield similar urine excretion (output). The sex-specific transporter patterns indicate different segmental transport loads between the sexes. Nonetheless, the question remains: *Why should the transporter pattern be different between the sexes?* The answer may lie, in part, in the paradoxical NHE3 transport capacity observed in the rat proximal tubule [18]: NHE3 expression is higher in female compared to male rats, but NHE3 also exhibits a higher degree of phosphorylation, which is a marker for localization to the base of the microvilli and lower NHE3 activity [27, 28]. We hypothesize that the lower baseline proximal tubule NHE3 activity in females may represent reserve capacity that can be utilized to significantly increase reabsorption when presented with glomerular hyperfiltration in order to limit natriuresis and diuresis. For example, during pregnancy and lactation both GFR and tubular transport are increased without a corresponding rise in blood pressure [29]. To investigate this hypothesis, we conduct simulations in which we raise GFR and determine urine output under three settings: (i) baseline transport parameters in the kidney of a man, (ii) baseline transport parameters in the kidney of a woman (which we refer to as the “non-adaptive female model”), and (iii) renal transport parameters in a woman, with the reserve NHE3 transport capacity activated (which we refer to as the “adaptive female model”).

Model simulations are limited to the consideration of glomerular hyperfiltration, tubular hypertrophy, and NHE3 activation; thus, these are by no means pregnancy/lactation simulations, even though some of the parameter choices are motivated by pregnancy-induced changes. In all three models, SNGFR is assumed to increase by 50% in superficial and 11% in juxtamedullary nephrons, consistent with GFR increases reported in pregnant women [30]. For the male and the non-adaptive female models, transport parameters are kept at baseline values (see [5] and Table 1). For the adaptive female model, we simulate the activation of reserve transport capacity by increasing NHE3 activity to 20% above male baseline value. Tubular hypertrophy is represented by increasing the model proximal tubule length by 20% [31]. No other pregnancy-related renal transporter changes are represented as we aimed to focus on the impact of hemodynamics and proximal tubule NHE3 by asking: *Given increases in GFR similar to those observed in pregnancy, to what extent would the higher filtered load be compensated for in the female kidney with the proposed pregnancy-induced morphological and transport changes along the proximal tubules? And how does the resulting solute and water handling compare to the male and non-pregnant female kidneys?*

Predicted segmental solute and water deliveries are shown in Fig. 4; segmental transport values are summarized in Fig. S3. First compare the male and non-adaptive female models. The lower NHE3 activity in the non-adaptive female model results in 8% higher Na^+^ delivery to the loops of Henle, compared to male. But the larger Na^+^ transport capacity along the female thick ascending limbs and distal segments more than compensates; consequently, urinary Na^+^ excretion is lower (22%) in the non-adaptive female model, compared to male (Fig. 4).

**Figure 4.**
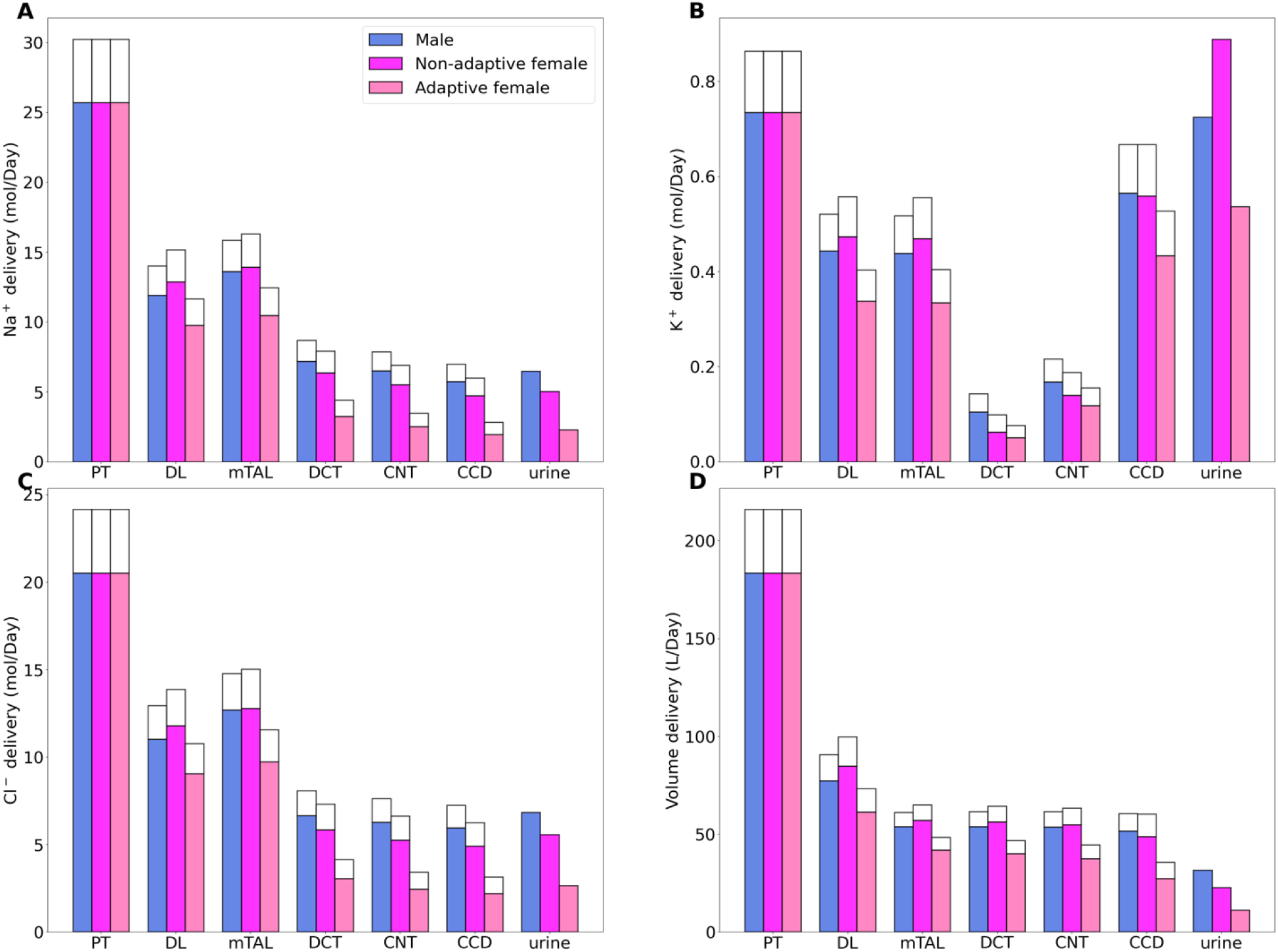
Effect of hemodynamic and PT changes on segmental delivery. Deliveries of key solutes (A–C) and fluid (D) to individual nephron segments, in male and female models 43% increase in GFR alone, and in the female model with 43% increase in GFR as well as NHE3 upregulation and PT hypertrophy and NHE3 upregulation. Delivery values are computed per kidney. Notations are analogous to Fig. 2.

Next, we consider the adaptive female model. The higher NHE3 activity and larger proximal transport area in the adaptive female kidney model results in a larger fraction of the filtered Na^+^ and K^+^ load being reabsorbed along the female proximal tubule compared to male (Fig. S3). The difference in Na^+^ flow (higher in the male model) is further augmented along downstream segments due to the higher (baseline) Na^+^ capacity along the thick ascending limbs in the female models. Eventually, that results in urinary Na^+^ excretion that is almost three times lower in women compared to men (Fig. 4).

Despite the higher fractional reabsorption, the elevated GFR induces severe natriuresis, diuresis, and kaliuresis in all three models. The predicted Na^+^ excretion is 17-, 38- and 47-fold higher than baseline in adaptive females, non-adaptive females and males, respectively. Because the reabsorption of Cl^-^ and water are largely coupled to Na^+^, similar trends are obtained for the Cl^-^ excretion and urine output (Fig. 4). The higher Na^+^ flow increases tubular K^+^ secretion in both sexes, resulting in markedly elevated K^+^ excretion that is comparable to filtered K^+^ load (Fig. 4B).

Taken together, these simulations indicate that tubular hypertrophy, activation of reserve NHE3 capacity, and the higher (baseline) distal transport capacity together can attenuate the excessive natriuresis, diuresis, and kaliuresis that is predicted to arise due to hemodynamics changes similar to those observed in pregnancy. However, to achieve baseline urine output and excretion, additional changes are required, including transport adaptation along the distal segments [32].

### Female renal transporter pattern may explain sex differences in response to a class of anti-hypertensive medications

Sex differences are reported in the prevalence and severity of cardiorenal diseases, and in pharmacodynamics [33, 34]. Of note is the angiotensin converting enzyme (ACE) inhibitors, a popular class of anti-hypertensive drugs target the renin-angiotensin system, which plays a key role in blood pressure regulation. ACE inhibitors lower blood pressure by reducing the production of angiotensin II (Ang II), a hormone that provokes vasoconstriction and fluid volume retention. ACE inhibitors are reported to be less effective in lowering blood pressure in women, compared to men [34–37]. Given that Ang II acts along proximal and distal tubular segments to increase Na^+^ transport, and thus, enhance salt and water retention, a reasonable question is: *To what extent can the lesser efficacy of ACE inhibitors’ in women be explained by the sex differences in renal transporters, especially along the distal segments?*

To answer that question, we simulate ACE inhibition in the two sexes by decreasing NHE3 activity along the proximal tubule by 25%, by decreasing NKCC2 activity along the thick ascending limbs by 60%, and by decreasing the activities of NCC, NaK-ATPase, ROMK, and ENaC along distal convoluted tubule by 60%. ACE inhibition affects the proximal tubule, thick ascending limbs and downstream segments, and we assume that SNGFR is unchanged [38, 39]. As noted above, because we assume lower NHE3 activity, proximal Na^+^ reabsorption is lower in women compared to men, resulting in higher NaCl and volume flow into the thick ascending limbs in women. Despite the higher Na^+^ delivery to the thick ascending limbs, the lower NKCC2 activity, following ACE inhibition, results in a small increase in Na^+^ reabsorption along the segment (~4%). In contrast, despite the reduction in NCC and ENaC activities along the distal segments, the higher tubular Na^+^ flow increases Na^+^ reabsorption, but to substantially different degrees in the two sexes. Due to their higher distal transport capacity, Na^+^ reabsorption increases by 52% in women, compared to only 31% in men. Similar effects are observed in water transport. Overall, ACE inhibition induces natriuresis and diuresis, but the increase in Na^+^ excretion is about half in women relative to men; similarly, urine output increase is about two-thirds lower in women. These simulations suggest that the blunted natriuretic and diuretic effects of ACE inhibition in women can be attributed, in part, to their higher distal baseline transport capacity.

ACE inhibition affects most nephron segments. To dissect the impact of proximal versus distal inhibition by ACE, we conducted “proximal inhibition” simulations in which NHE3 activity was inhibited along the proximal tubule, and “distal inhibition” simulations in which NKCC2, NCC, NaK-ATPase, ROMK, and ENaC were inhibited along the downstream segments. Segmental delivery and transport of Na^+^, K^+^, Cl^-^, and fluid are shown in Figs. S4 and S5. Simulation results indicate a larger effect, in terms of urinary excretion, from proximal inhibition compared to distal.

## Discussion

The present study seeks to answer the question: What is the physiological implication of our postulated differences in renal membrane transporter distribution between men and women? Given that transport activity pattern is mostly uncharacterized in the human kidney, let alone the sex differences, we extrapolate our knowledge from rodents to humans. In the first modeling study of nephron transport in humans [5], we demonstrated that a model that combines known renal hemodynamics and morphological parameters in humans, with membrane solute and water transport parameters similar to rats, can predict segmental deliveries and urinary excretion of volume and key solutes that are consistent with human data. Transport parameters used in that study were based on the male rat. How might the results be different if we had modelled a woman by adopting transport parameters from the female rat [18]?

In rats, baseline Na^+^ transport of the proximal nephron is much lower in females, due to the significantly lower NHE3 activity and smaller tubular size. However, GFR and thus filtered load are also lower in females. In contrast, GFR is similar between men and women, as are their kidney size. Hence, in humans we focus on the renal transport pattern as the major contributor to any sex differences in nephron transport By assuming sexual dimorphisms similar to that in rats [18], model simulations indicate that the lower fractional reabsorption of filtered Na^+^ and water along the proximal nephron in women can be compensated by more abundant transporters as well as the increased solute delivery to these transporters in downstream tubular segments (Fig. 2), so that urine output and excretion rates are similar in men and women. It should be noted that the main proximal tubule sodium reabsorptive pathway, NHE3, is a Na^+^/H^+^ exchanger which contributes to acid excretion bicarbonate reabsorption and NH4^+^ secretion. How sex differences in NHE3 and other transporters impact acid-base balance is a matter of current investigation by a number of labs [40]. While it is reassuring that both the male and female renal transporter patterns can match urinary excretion to dietary intake, one may wonder why should the pattern be different between men and women in the first place? A notable sex difference observed in rats, that we assume translates to humans, is that NHE3 is actually more abundant along the proximal tubules in females, but a larger fraction of NHE3 is phosphorylated and thus inactive. In other words, females appear to have a larger proximal Na^+^ transport reserve. Additionally, Na^+^ transport along the distal nephron is exquisitely regulated, and its larger contribution to total renal Na^+^ transport in females may also better prepare them for the fluid retention adaptations required during pregnancy and lactation.

Pregnancy involves a remarkable orchestration of changes in the majority of the physiological systems, including essentially all aspects of kidney function. Mild increases in proteinuria, glucosuria, lower serum osmolality, and reductions in serum Na^+^ levels are often reported in pregnancy [41–43]. Renal and systemic hemodynamics are both drastically altered by the marked volume expansion and vasodilation. As such, renal plasma flow increases by up to 80% and GFR increases by 50% [44]. Does the female renal transporter pattern better prepare women to handle the much larger filtered load? Model simulations indicate that by dephosphorylating and thereby activating the reserve NHE3 in the proximal nephron, the female kidney can limit excessive natriuresis and diuresis, compared to males (see Fig. 4) and also indicate that the larger transport capacity in the distal nephron of the female kidney renders women better equipped to handle (i.e., retain a significantly more of) the larger filtered load in pregnancy. Nevertheless, to completely compensate for the heightened load in pregnancy, further alternations in tubular transport are required [32, 45, 46].

The kidney plays an indispensable role in blood pressure regulation [47]. To maintain arterial circulation and blood pressure, the kidney tightly controls its own renal perfusion pressure, renal perfusion and extracellular volume. Renal artery perfusion pressure directly regulates sodium excretion, in a process known as pressure natriuresis, and influences the activity of various vasoactive systems such as the renin–angiotensin system (RAS). Major differences are evident between men and women in the RAS that may manifest in differences in blood pressure regulation [37, 48]. Indeed, the prevalence of hypertension is higher in men compared with women prior to the onset of menopause.

Historically, most epidemiological and experimental data collected on the progression of hypertension and chronic renal disease were collected in men, as were data on the effectiveness of therapeutic interventions. Few clinical trials examining the health benefits of RAS inhibition have reported data for both sexes separately, even when women have been included. Falconnet et al. reported that, following a 20-day treatment with lisinopril at 20 mg/day, men responded with a larger decrease in ambulatory blood pressure compared with women [34]. Their data suggest that over time, the effectiveness of ACE inhibition is lessened in women. That conclusion is supported by additional cardiovascular studies in patients with congestive heart failure and following myocardial infarction in which ACE inhibition confers less cardiovascular benefit to women compared with men as assessed by total mortality [36]. *To which extent can the sex differences in the response to ACE inhibitors be attributable to sex differences in renal tubular transport?* To answer this question, we simulated kidney function following ACE inhibition, and compared the predicted natriuretic and diuretic responses in men and women. We assumed that ACE inhibition generates the same fractional reduction in key Na^+^ transporter activities in men and women. With this key assumption, model simulations suggest that the higher distal transport capacity in women results in greater attenuation in natriuresis and diuresis, compared to men (Fig. 5), which, when taken in isolation, may result in a small reduction in blood pressure. In male rodents, angiotensin II stimulates and ACE inhibitors attenuates proximal tubule Na^+^ reabsorption through actions on NHE3 and Na,K-ATPase [49, 50]. While similar studies in females are, to our knowledge, lacking, the lesser sodium reabsorption in female versus male rats indicates that ACE inhibitors would have more proximal sodium transport to inhibit in males than in females. While these results suggest that sex differences in tubular transport may contribute to the lesser efficacy of ACE inhibitors in women, one must not neglect the other systems that are affected by or interact with the action of ACE inhibitors [51].

**Figure 5.**
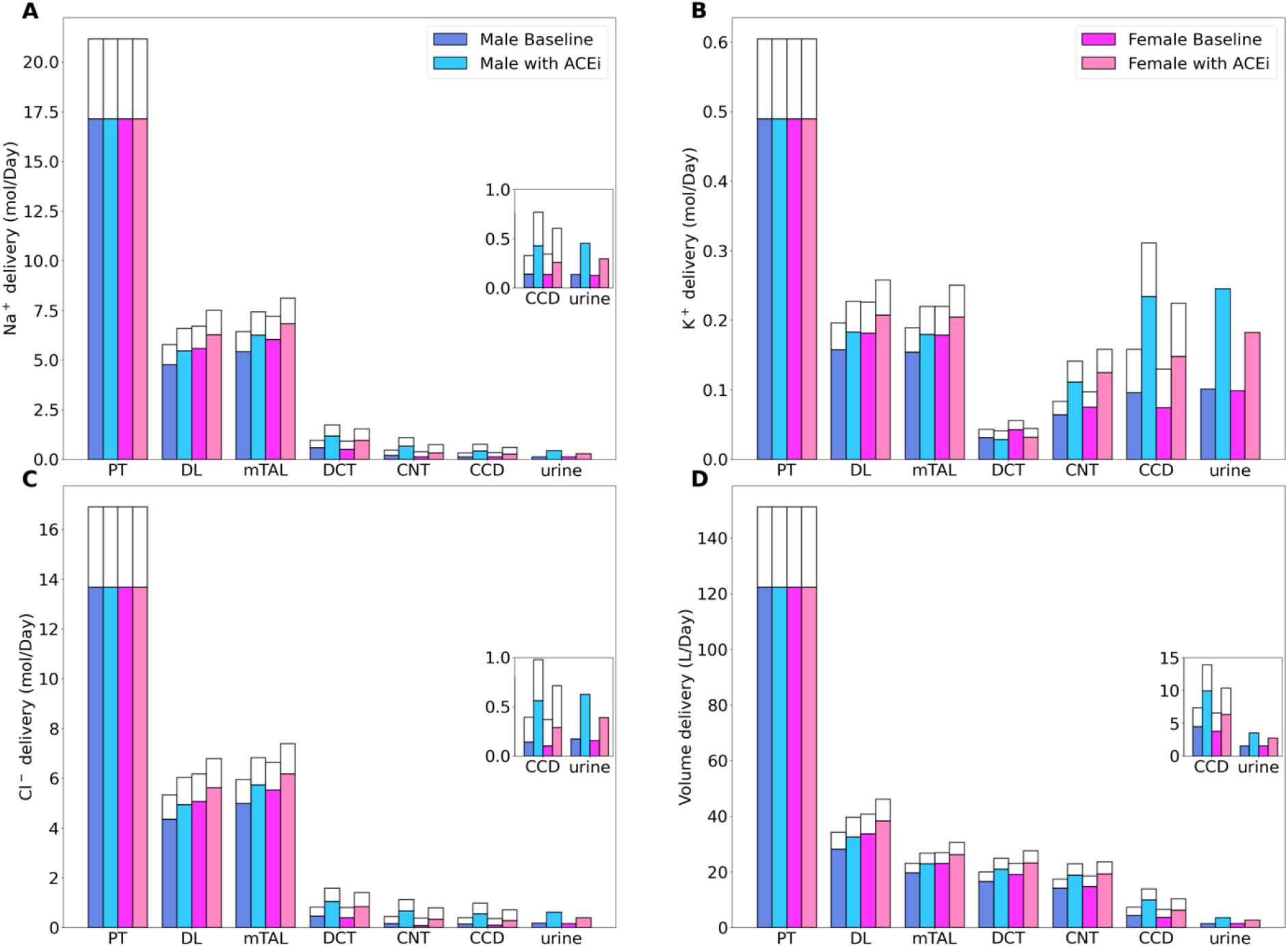
Comparison of solute delivery (*A*–*C*) and fluid delivery (*D*) obtained for base case and for ACE inhibition for both sexes. Delivery values are computed per kidney. Notations are analogous to Fig. 2.

In our simulations, we assume that ACE inhibition has the same effect on kidney transport in men and women. However, major sex differences in the RAS have been identified, including how much substrate is produced and how angiotensin interacts with receptors. For example, in both rats and humans, females have greater circulating levels of AGT [52], causing an overall greater amount of angiotensin to be flowing through the system. Also, male rats have been shown to have greater Ang II levels [53] while female rats have greater levels of Ang (1–7) [54]. However, a greater level of ACE2 activity (which converts Ang II to Ang (1–7)) has been measured in male rats [54]. Indeed, males and females exhibit differing responses to Ang II. Changes in Ang II levels produce smaller changes in blood pressure in female rats [52, 53]. That discrepancy may be attributed to differences in receptor expression, inasmuch as male rats have greater AT1R expression and lower AT2R expression than female rats [53–55]. Estrogen also reduces the half-life of AT1R-bound Ang II [56]. Given the above observations, it is likely that ACE inhibitors affect nephron transport differently in men and women, but those effects have not been sufficiently well characterized.

In sum, we have extended our work in sex-specific modeling of kidney function in the rat [19–21, 57, 58] to human. Compared to rodents, human kidneys are more similar between the sexes, in terms of morphology and hemodynamics. Nonetheless, our study suggests that sexual dimorphism in renal transport pattern similar to that reported in rodent may better prepare women for the heightened demands on the kidney during pregnancy, and may also contribute to the two sexes’ differential response to ACE inhibition. The present sex-specific nephron models can be used as essential components in more comprehensive models of integrated kidney function. For instance, computational models of whole-body blood pressure regulation typically include a highly simplified representation of the kidney (e.g., [59, 60]). Incorporating elaborate sex-specific nephron transport models would allow more accurate simulations of the very commonly prescribed anti-hypertensive treatments that target the kidney (e.g., ACE inhibitors, diuretics) and analysis of any sex differences in drug efficacy. The present model can be combined with models of renal oxygenation that represent blood flow and tubular metabolism [61–65]. The resulting comprehensive sex-specific models of kidney oxygenation would allow one to conduct a comprehensive *in silico* study of factors that give rise to the impact of sex on the progression of chronic kidney disease [66].

It has become increasingly clear that experimental and clinical results obtained in one sex cannot be extrapolated to both sexes without sufficient justification. A better understanding of the sex differences in renal physiology and pathophysiology is a necessary step toward a true understanding of the complicated transport and excretion processes in the kidney, and, if needed, toward sex-specific and more effective therapeutic treatments for men and women.

## Materials and Methods

We previously developed an epithelial cell-based model of solute transport along a superficial nephron of a human (male) kidney. In this study, we extend that model (i) to include the juxtamedullary nephron populations, and (ii) to represent nephron populations in a woman’s kidney. (Superficial nephrons turn at the outer-inner medullary boundary, whereas juxtamedullary nephrons reach into differing levels of the inner medulla.) The model represents six classes of nephrons: a superficial nephron (denoted “SF”) and five juxtamedullary nephrons (denoted “JM”). The superficial nephrons account for 85% of the nephron population, and extend from the Bowman’s capsule to the papillary tip. The remaining 15% of the nephrons are juxtamedullary that possess loops of Henle that reach to different depths in the inner medulla; most of the long loops turn within the upper inner medulla. The model nephron is represented as a tubule lined by a layer of epithelial cells, with apical and basolateral transporters that vary according to cell type. The model assumes that the connecting tubules coalesce successively within the cortex, resulting in a ratio of loop-to-cortical collecting duct of 10:1 [67]. In the inner medulla, the collecting ducts again coalesce successively. A schematic diagram for the model is shown in Fig. 1. Baseline single-nephron glomerular filtration rate (SNGFR) is set to 100 and 133 nl/min for the superficial and juxtamedullary nephrons, respectively, in both men and women. Assuming a total of 1 million nephrons in each kidney, this yields a single-kidney GFR of 105 mL/min.

The model accounts for the following 15 solutes: Na^+^, K^+^, Cl^-^, HCO_3_^-^, H_2_CO_3_, CO_2_, NH_3_, NH_4_^+^, HPO_4_^2-^, H_2_PO_4_^-^, H^+^, HCO_2_^-^, H_2_CO_2_, urea, and glucose. The model is formulated for steady state and consists of a large system of coupled ordinary differential equations and algebraic equations (see model equations below). Model solution describes luminal fluid flow, hydrostatic pressure, luminal fluid solute concentrations and, with the exception of the descending limb segment, cytosolic solute concentrations, membrane potential, and transcellular and paracellular fluxes.

In this study, we develop an analogous multi-nephron model for a woman. We follow the approach implemented in our recent studies [19, 20] in which we formulated sex-specific epithelial cell-based models of solute transport along the nephrons of the male and female rat kidneys. The male and female multi-nephron models presented here account for sex differences in the expression levels of apical and basolateral transporters [18, 68].

Key model equations are described in the supporting information. Model parameters can be found in Ref. [5]. Key differences in nephron transport parameters between men and women are summarized in Table 1. The model is implemented in Python and can be accessed via GitHub.

## Acknowledgement

This research was supported by the Canada 150 Research Chair program, by the Natural Sciences and Engineering Research Council of Canada, via a Discovery award to ATL, and by the National Institutes of Health NIDDK, via 2R01DK083785 and R56DK123780 to AAMcD.

## Supporting Information

S1 Text. Model equations and supplemental figures.

